# *In vivo* cross-linking and transmembrane helix dynamics support a bidirectional non-piston model of signaling within *E. coli* EnvZ

**DOI:** 10.1101/206888

**Authors:** Rahmi Yusuf, Tuyết Linh Nguyễn, Annika Heininger, Robert J. Lawrence, Benjamin A. Hall, Roger R. Draheim

**Affiliations:** School of Pharmacy and Biomedical Sciences, University of Portsmouth, Portsmouth, PO1 2DT, England, UK; Institute of Biochemistry, Biocenter, Goethe University Frankfurt, D-60438 Frankfurt, Germany; MRC Cancer Unit, University of Cambridge, Cambridge, CB2 0XZ, England, UK; Institute of Biological and Biomedical Science, University of Portsmouth, Portsmouth, PO1 2DT, England, UK

**Keywords:** porin balance, transmembrane communication, coarse-grained molecular dynamics, sulfhydryl99 reactivity, transmembrane helix dynamics

## Abstract

In Gram-negative bacteria, porins span the outer membrane and control the influx of several prominent groups of antibiotics. Thus, it should not be surprising that expression of these porins is often altered in clinical isolates exhibiting multidrug resistance (MDR). The major regulator of porin expression in *Escherichia coli* is EnvZ, a canonical sensor histidine kinase (SHK). It allosterically processes periplasmic interactions with MzrA and cytoplasmic osmosensing into a single unified change in the ratio of its kinase and phosphatase activities. Unfortunately, the role of the EnvZ transmembrane domain (TMD) in bidirectional communication of these signals remains not well understood. Here, we employed *in vivo* sulfhydryl-reactivity to probe the dynamics of the TM2 helices and demonstrate that upon stimulus perception, only the region proximal to the periplasm undergoes conformational rearrangement. Furthermore, *in silico* coarse-grained molecular dynamics (CG-MD) simulations with aromatically tuned variants of EnvZ TM2 demonstrate the existence of both tilting and azimuthal rotational components to transmembrane communication while ruling out piston-type repositioning of TM2. Finally, in contrast to a similar analysis of TM1, we identified position-specific mutants possessing a “flipped” phenotype by dual-color fluorescent reporter analysis suggesting that both the periplasmic and cytoplasmic ends of TM2 are critical for maintenance of EnvZ signal output. Taken together, these data strongly support that EnvZ employs a non-piston-type mechanism during transmembrane communication. We conclude by discussing these results within the context of allosteric processing by EnvZ and propose that these results can be used to predict and classify transmembrane communication by various SHKs.

**Importance:** The EnvZ sensor histidine kinase serves as the major regulator of porin expression within *Escherichia coli*. A long-standing question is how stimulus perception by a bacterial receptor on one side of a biological membrane is transmitted to the opposite side of the membrane. To address this question, we monitored the dynamics of the transmembrane domain of EnvZ *in vivo* and coupled these results with *in silico* simulations of membrane-embedded EnvZ transmembrane domains. Taken together, these results demonstrate that detection of osmotic stress by the cytoplasmic domain of EnvZ results in non-piston communication across the inner membrane of *E. coli.* Thus, in addition to understanding how EnvZ regulates porin balance and antibiotic influx, these results contribute to answering the long-standing question of how transmembrane communication is performed by bacterial receptors. Our work concludes with a framework that correlates receptor domain composition and signal transduction mechanisms that could be employed by other research groups on their particular receptors of interest.

## Introduction

Most porins involved in antibiotic transport by Gram-negative bacteria belong to the classical OmpF and OmpC families (1). Transcription of these porins is governed by the intracellular concentration of phospho-OmpR (OmpR-P), which is controlled by EnvZ in response to changes in periplasmic interactions with MzrA and environmental osmolarity (2-7) (Fig. 1A). At low intracellular levels of OmpR-P, transcription of *ompF* is upregulated, whereas at higher levels of OmpR-P, transcription of *ompF* is repressed and transcription of *ompC* is activated. This results in a predominance of OmpF at low osmolarity and OmpC at higher osmolarities or in the presence of MzrA (8-10) (Fig. 1B). Dramatic modification of porin balance, which has been observed within clinical isolates from patients undergoing antibiotic treatment, strongly supports further characterization of the underlying mechanisms of porin regulation by EnvZ (11-17). In addition, it was-recently shown that mutations in EnvZ within a porin-deficient (*ompC^-^ ompF^-^*) *E. coli* strain resulted in increased carbapenem resistance (18). Thus, in addition to maintenance of porin balance, EnvZ also plays another not well-understood role in □mediating antibacterial resistance that warrants further elucidation.□

**Figure 1.**
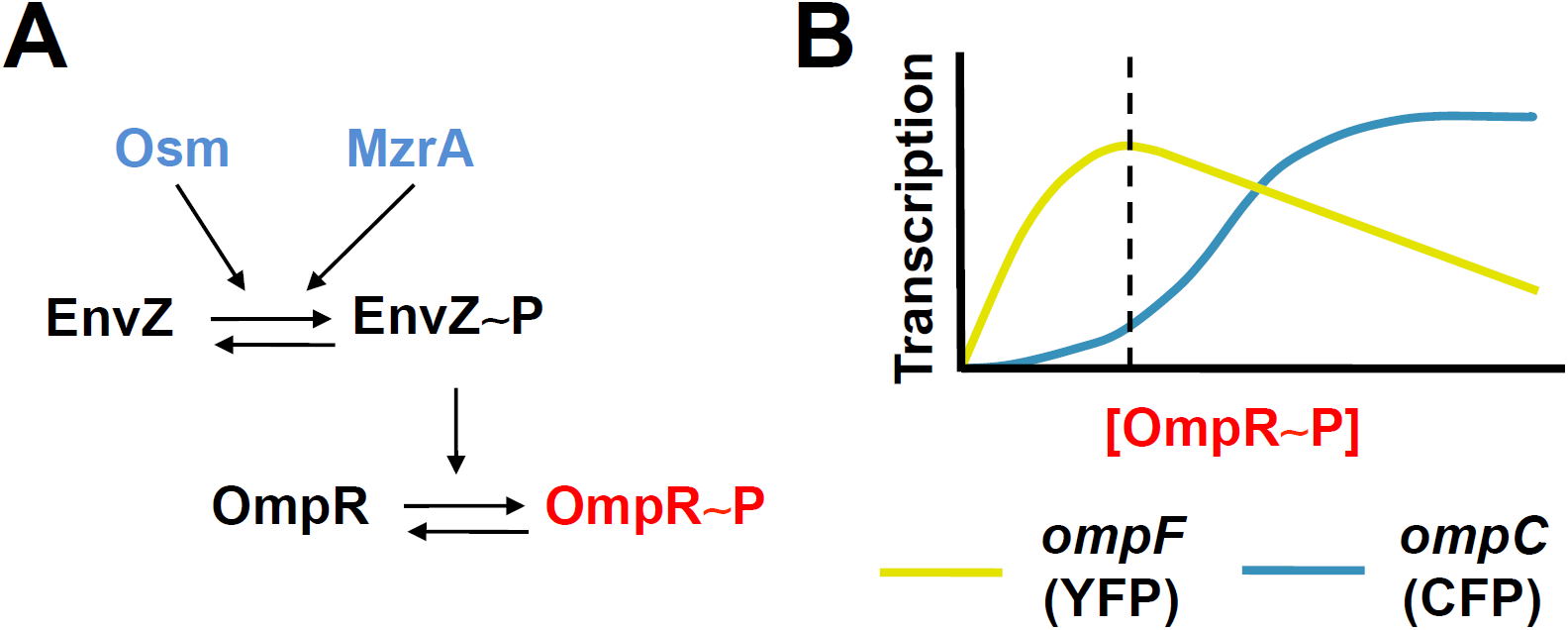
Monitoring modulation of EnvZ signal output upon stimulus perception. (A) EnvZ signal output controls porin expression. It is a bifucntional sensor histidine kinase with both kinase and phosphatase activities. The ratio of these activities is modulated by the presence of extracellular osmolarity and the absence/presence of MzrA (blue). (B) The intracellular level of phosphorylated OmpR (OmpR-P) is controlled by EnvZ signal output and OmpR-P levels govern transcription of *ompF* and *ompC*. Strains MDG147 and EPB30 (Δ*envZ*) contain transcriptional fusions of *yfp* to *ompF* (yellow) and *cfp* to *ompC* (cyan), which facilitates easy monitoring of intracellular OmpR-P levels by measuring the CFP/YFP ratio. The dashed line indicates an estimation of the baseline OmpR-P levels from EPB30/pRD400 cells grown under the low-osmolarity regime (0% sucrose).

A substantial body of evidence suggests that EnvZ participates in bidirectional allosteric processing. A cytoplasmic fragment of EnvZ (EnvZ_c_), consisting of residues Arg-180 through Gly-450, has been shown to mediate physiologically appropriate responses to increasing NaCl and sucrose concentrations *in vitro* and to increasing sucrose *in vivo* (19). These results led the suggestion that the peripalsmic and transmembrane domains of EnvZ play a negligible role in osmosensing with the exception of anchoring EnvZ within the membrane and thereby reducing a three-dimensional search by OmpR to a two-dimensional search (19). In addition, it was initially suggested membrane anchoring might reduce the conformational dynamics of EnvZ to physiologically relevant levels, as EnvZ_c_ was found to have greater specific activity than full-length membrane-bound EnvZ (19, 20). Further studies demonstrated that peripheral interactions between the cytoplasmic domains and the lipid membrane were essential in increasing the kinase activity of the cytoplasmic domain of EnvZ, including the observation of a roughly 25-fold in EnvZ_c_ signal output while maintaining regulation, *i.e.* a 2-fold increase in autophosphorylation, in response to sucrose *in vitro* (21). In summary, these results confirm that molecular dynamics within the cytoplasmic domain of EnvZ govern response to osmolarity and that lipid-receptor interactions play a modulating the signal output of EnvZ.

Alternatively, MzrA, <u>modulator</u> of <u>EnvZ</u> and <u>OmpR</u> protein <u>A</u>, was identified as a suppressor of a Δ*bamB* Δ*degP* double mutation. Upon further analysis, it was shown that MzrA is an upstream regulator of EnvZ signal output that is independent of signal output modulation due to osmolarity, pH and procaine (6). MzrA was subsequently shown to localize to the inner membrane and interact with EnvZ within the peripalsmic space (7) to serve as a connector between the CpxA/CpxR and EnvZ/OmpR signaling circuits (6). It is also known as a connector of which only a few examples have been found (22-24).

When taken together, these results demonstrate that the TMD of EnvZ is responsible for allosterically coupling sensory input from the attached periplasmic and cytoplasmic domains. By extension, bidirectional communication through the TMD is required in order to properly modulate porin expression. Thus, we were interested in how cytoplasmic osmosensing could modulate the conformational dynamics of the TMD and subsequently that of the periplasmic domain. Understanding how EnvZ transduces signal across the biological membrane would be a significant step toward direct manipulation of porin balance in bacterial cells exhibiting MDR.

To address these lines of enquiry, we have created a single-Cys-containing library of EnvZ mutants across the second TM helix (TM2) that physically connects the periplasmic and the cytoplasmic domains of EnvZ. *In vivo* analysis of this library revealed three regions within the TM domain, of which the periplasmic region undergoes a non-piston-type conformational change. CG-MD simulations of the aromatically tuned and wild-type EnvZ TM2 variants supported these results. We conclude by discussing these results in the context of governance of porin balance and that mechanisms of transmembrane communication can be classified based on the periplasmic domains that the bacterial receptor possesses.

## Results

### Creating a single-Cys-containing library within TM2 of EnvZ

We previously created a Cys-less version of EnvZ from *E. coli* that had its sole Cys residue changed to an Ala residue (C277A). The Cys-less variant is expressed from pRD400, which results in the addition of a seven-residue linker (GGSSAAG) and a C-terminal V5 epitope (GKPIPNPLLGLDST). We previously found that the Cys-less version of EnvZ had similar steady-state signal output and response to environmental osmolarity as the wild-type version of EnvZ making it suitable for comparisons of *in vivo* sulfhydryl-reactivity and signal output analysis. We initially determined that no major rearrangements occur along the TM1-TM1’ interface upon stimulus perception (25). As minimal change was observed along this helical interface in response to osmolarity, we continued by examining the TM2-TM2’ interface. We began by determining which residues comprise TM2 by subjecting the full EnvZ sequence to DGpred (26) and TMHMM v2.0 (27), which suggested that Leu-160 to Ile-181 and Leu-160 to Ile-179 comprise TM2 respectively. Based on these analyses, we employed site-directed mutagenesis using the Cys-less variant as a template to create a library of single-Cys-containing EnvZ proteins that spanned from positions 156 to 184 (Fig. 2). We observed that nearly the entire library was expressed within EPB30/pRD400 cells grown under the low‐ or high-osmolarity regime. Variants possessing a Cys at position 156 showed low levels of expression when grown under the low-osmolarity (0% sucrose) regime. However, when grown under the high-osmolarity regime, no variants showed reduced expression level (Fig. S1). These results indicate that the library was suitable for further *in vivo* experimentation.

**Figure 2.**
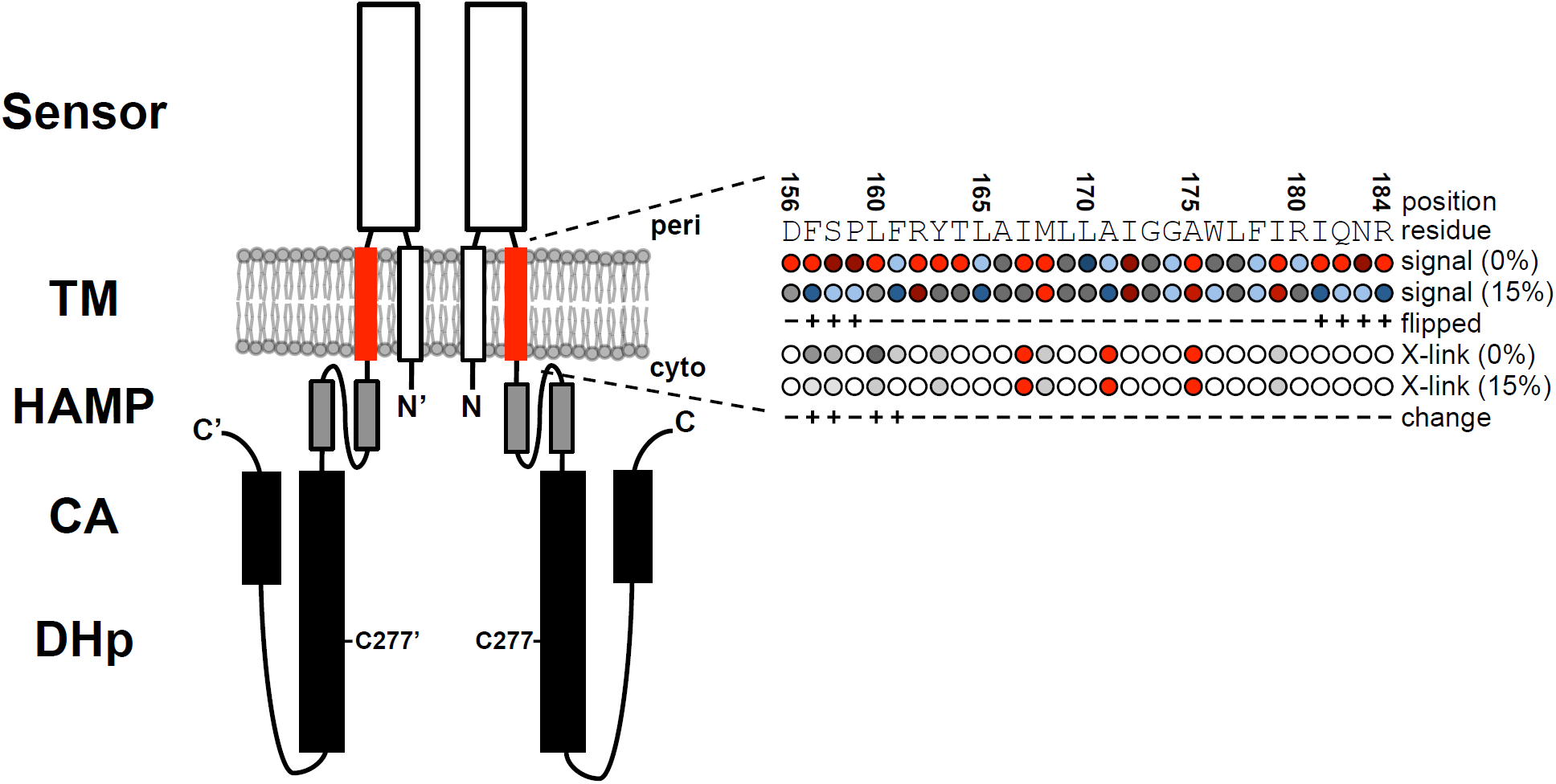
EnvZ functions as a homodimer with a cytoplasmic N-terminus, the first transmembrane helix (TM1, white), a large periplasmic domain (sensor, white), the second transmembrane helix (TM2, red), a membrane-adjacent HAMP domain (grey) and the cytoplasmic domains responsible for dimerization and histidylphosphortransfer (DHp, black) and catalytic ATPase activity (CA, black). The position of the original Cys-277 residue that was mutated to Ala to produce the Cys-less EnvZ is provided. The residues subjected to Cys substitution and their position in the primary sequence is provided. Signal output from each single-Cys-containing variant is compared to the Cys-less (C277A) variant: less than 50% of Cys-less (light blue), between 50% and 75% of Cys-less (dark blue), between 75% and 125% of Cys-less (grey), between 125% and 200% (dark red) and greater than 200% (light red). Residue positions exhibited flipped signal output are indicated with a plus. The extent of sulphydryl-reactivity is also presented in five categories based on dimer-to-monomer ratio: no dimer present (white), less than 0.05 (light grey), between 0.05 and 0.2 (medium grey), between 0.2 and 0.5 (dark grey) and greater than 0.5 (red). Positions that exhibit a significant change in cross-linking between the low‐ and high-osmolarity regimes are indicated with a plus.

### Mapping TM2 surfaces important for maintenance of EnvZ signal output

We began by expressing each of the single-Cys-containing variants in EPB30/pRD400 cells, which allowed us to measure CFP fluorescence, YFP fluorescence, and to calculate the CFP/YFP ratio that estimates steady-state EnvZ signal output (Fig. 1B). Cells expressing the Cys-less C277A were used as a baseline comparison (Fig. S2). When EPB30/pRD400 cells are grown under the low-osmolarity regime, a shift in signalling output toward the “on” or kinase-dominant state results in increased CFP fluorescence, reduced YFP fluorescence and an increase in the overall CFP/YFP ratio, while a shift toward the “off” or phosphatase-dominant state appears as decreased CFP, increased YFP and a decrease in CFP/YFP ratio (Fig. 3 and S3).

Several trends were observed during analysis of the library of Cys-containing EnvZ receptors. When EPB30/pRD400 cells were grown under the low-osmolarity regime, EnvZ was less tolerant of Cys substitutions at the N‐ and C-terminal regions of the library. At the N-terminus, signal output from receptors containing a Cys at positions 156, 162 and 163 were very elevated, exhibiting greater than a 5-fold increase in CFP/YFP, while receptors possessing a Cys in the C-terminus at positions 179, 181, 182 and 184 were elevated, possessing over a 2-fold increase in CFP/YFP. These boundary regions appear to flank a core of alternating increases and decreases in EnvZ signal output, as observed between residue positions 165 and 180, suggesting that multiple tightly packed EnvZ helices exist within the hydrophobic core of the inner membrane (Fig. 3A and 3B). When grown under the high-osmolarity regime, a pattern appeared where Cys substitutions resulted in significant decreases in signal output (Fig. 3C). Of the 29 mutants analysed, 13 supported less than 75% of the normal wild-type signal output.

**Figure 3.**
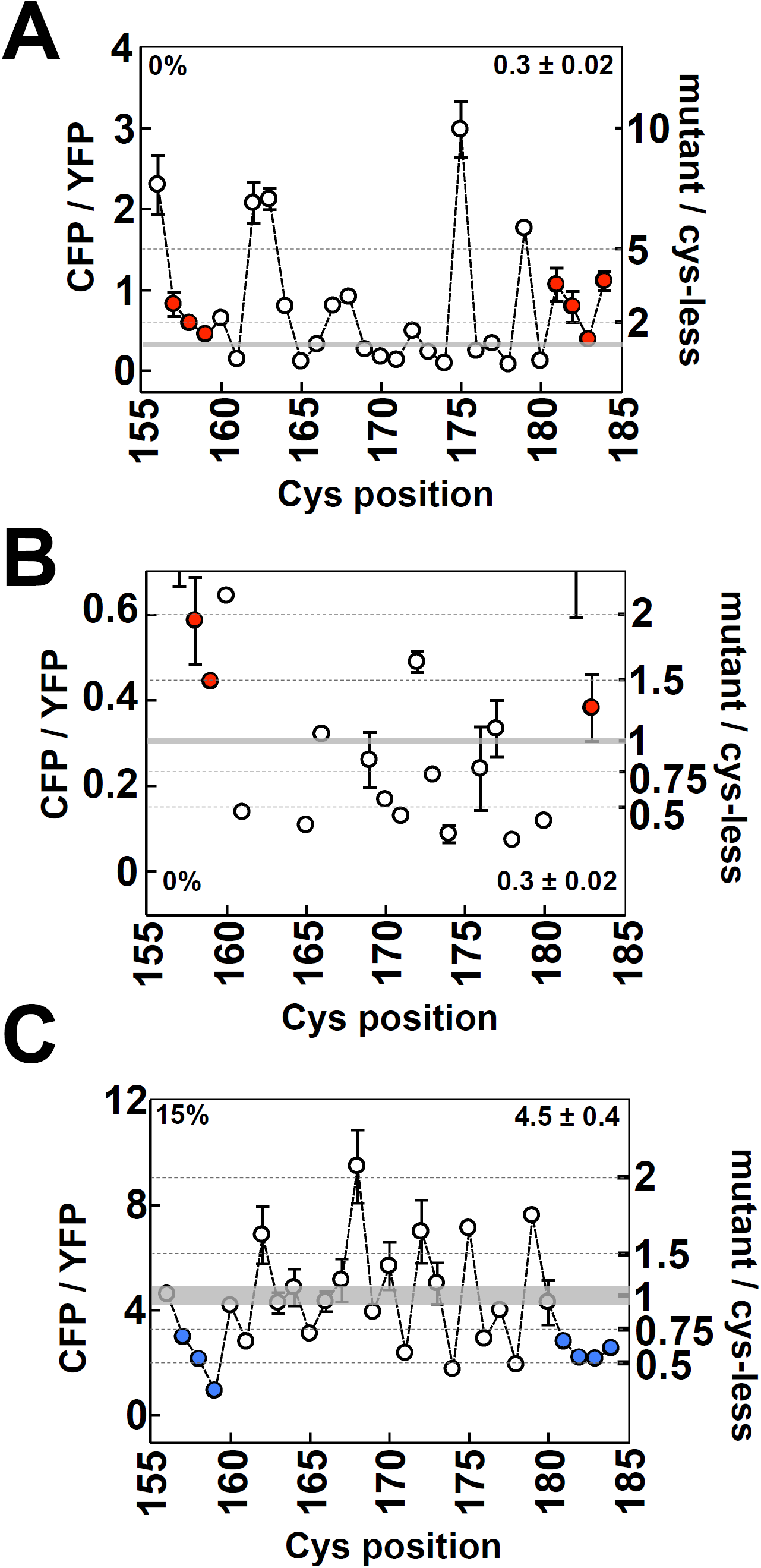
Signal output from the library of single-Cys EnvZ variants. (A) CFP/YFP from EPB30/pRD400 cells expressing one of single-Cys variants grown under the low-osmolarity (0% sucrose) regime. On the right axis, these CFP/YFP ratios are compared to the Cys-less (C277A) variant. (B) Magnified version of panel A in order to emphasise the region up to a 2-fold increase in CFP/YFP. (C) CFP/YFP from EPB30/pRD400 cells expressing one of the single-Cys variants grown under the high-osmolarity (15% sucrose) regime. On the right axis, these CFP/YFP ratios are compared to the Cys-less (C277A) variant. The flipped mutants are highlighted with a red dot in panel A (increased signal output) and a blue dot in panel C (decreased signal output). The shaded areas represent the mean signal output from the Cys-less variant of EnvZ with a range of one standard error of mean. These values are provided to aid in comparison. Error bars represent standard deviation of the mean with a sample size of n ≥ 3.

Most striking is the inverse effect on EnvZ signal output of the Cys substitutions that flank the hydrophobic core of TM2. For these residues, when grown under the low-osmolarity regime, the presence of a Cys resulted in an increase in signal output of more than 25%, i.e. shifted toward the on or kinase-dominant state (red dots in Fig. 3A) and a reduction in signal output of more than 25%, i.e. shifted toward the off or phosphate-dominant state, when grown under the high-osmolarity regime (blue dots in Fig. 3C). These flipped positions reside at the N‐ and C-terminal ends of the region we examined and outside of the proposed hydrophobic TMD core (Fig. 3). Within the hydrophobic core, Cys substitutions show similar changes when cells are grown under the low‐ and high-osmolality regimes. These results suggest that the flanking regions are not simply rigid structural conduits for signal transduction but may have higher-order roles in signal transduction, such as MzrA interaction or functioning as a control cable at the N‐ and C-terminal regions respectively (6, 7, 28-31).

### Identifying surfaces involved in TM2-TM2’ interactions

Sulfhydryl-reactivity experimentation is well-characterised and has been employed on many soluble and membrane-spanning proteins and higher-order complexes (32). The *in vivo* nature of this assay facilitated mapping of the TM2-TM2’ interface under different osmotic conditions, which is an important first step toward understanding how EnvZ processes allosteric inputs from periplasmic MzrA binding and cytoplasmic osmosensing into a single uniform modulation of bacterial porin balance (Fig.1). In a similar manner to mapping TM1-TM1’ interactions (25), Cys-containing EnvZ variants were expressed in EPB30/pRD400 cells and upon entering the early exponential phase (OD_600nm_ ≈ 0.25) they were subjected to 250 μM molecular iodine for 10 minutes analysed by non-reducing SDS-PAGE and immunoblotting (Fig. S4).

We observed three distinct regions within TM2. The N-terminal region (region I in Figure 3), comprised of residues 156 to 163, exhibited significant cross-linking under the low-osmolarity regime (0% sucrose) and almost no crosslinking under the high-osmolarity (15% sucrose) regime. The second region (II) consisting of positions 164 to 179, demonstrated altering low and high levels of disulphide-formation consistent the crossing of TM2 and TM2’ within the hydrophobic core of the TMD. The final region (III), from residues 180 to 184, shows no crosslinking (Fig. 4). It should be noted that this significant difference at the periplasmic end of the TMD between cells grown under the low‐ and high‐osmolarity regime was not observed during similar analyses of TM1 (25).

**Figure 4.**
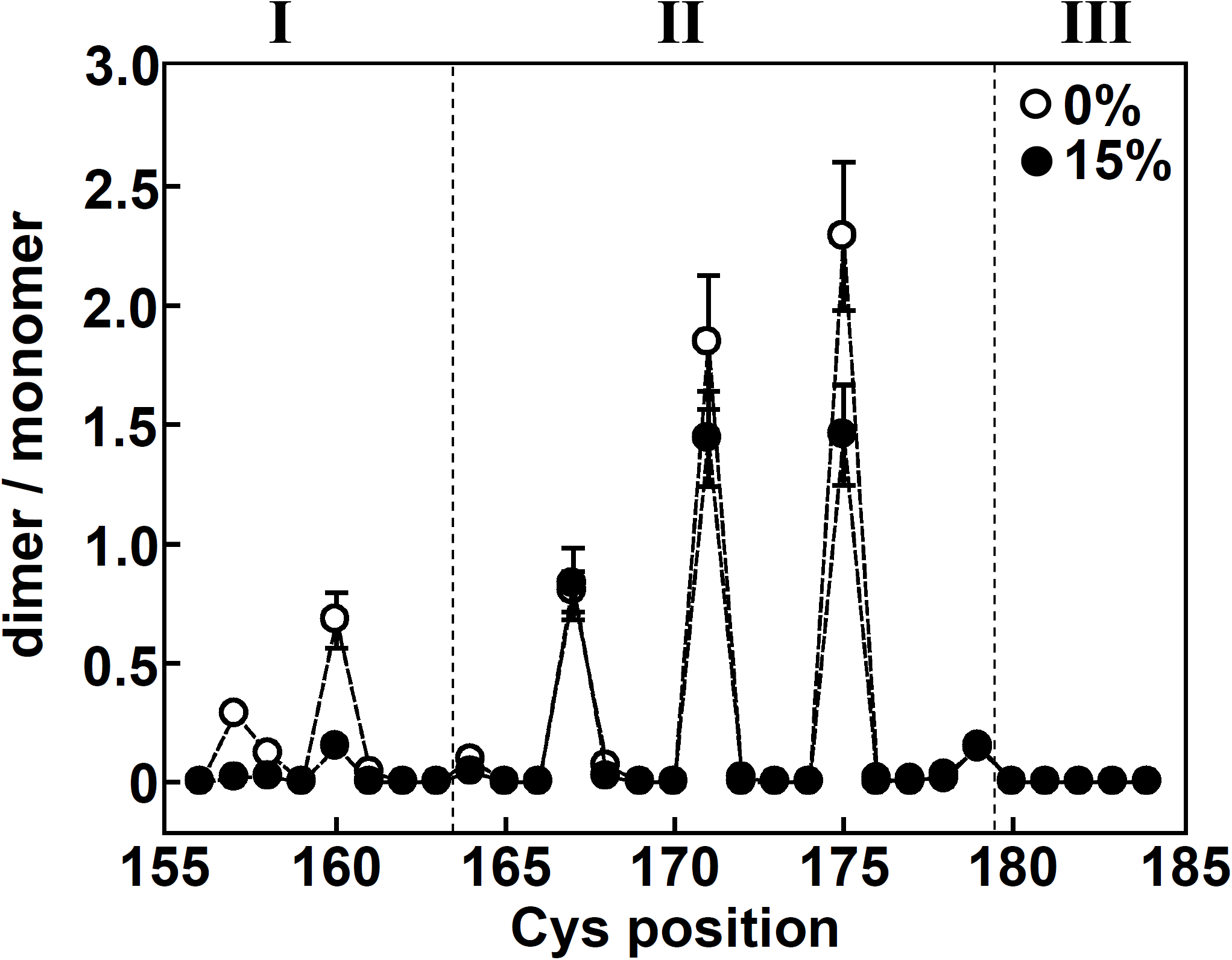
Extent of sulfhydryl-reactivity for each single-Cys-containing EnvZ variant. EPB30/pRD400 cells were grown under the low‐(empty circles, 0% sucrose) or high-osmolarity (filled circles, 15% sucrose) regimes and subjected to 250 μM molecular iodine for 10 minutes when their OD_600nm_ reached approximately 0.25. As shown in Figure S5, this allowed us to determine the dimer/monomer ratio represented on the Y-axis. Three distinct regions, denoted I, II and III were observed and are described in the text. Error bars represent the standard error of the mean with a sample size of n ≥ 3.

### Molecular simulations of the wild-type and aromatically tuned variants of EnvZ TM2

The piston-type model of transmembrane communication is founded upon the central tenant that the vertical position of TM2 relative to the lipid bilayer changes upon stimulus perception. Previous work with TM2 of Tar demonstrates that repositioning the aromatic residues at the cytoplasmic end of TM2, known as aromatic tuning (33), repositions the helix within a biological membrane (34) and that this repositioning causes an incremental change in signal output (33). These results served as an experimental framework to optimise SIDEKICK software capable of high-throughput parallelised coarse-grained molecular dynamics (CG-MD) simulations, which demonstrated that aromatic tuning repositioned the TM2 helix *in silico* in a manner consistent with both the *in vivo* results and a piston-type mechanism of transmembrane communication (35).

Based on these extensive results and the recent suggestion that mechanisms employed by SHKs during TM communication correlate with the presence of a membrane-adjacent HAMP domain (36), we performed analogous *in silico* experimentation with EnvZ, which possesses a membrane-adjacent HAMP domain but substitutes a periplasmic PDC/CACHE domain for the four-helix bundle present in Tar and NarQ (36-39). We previously performed aromatic tuning with TM2 of EnvZ and found that signal output was not correlated with the absolute vertical position of the aromatic residues as was shown with Tar TM2, which suggested that EnvZ does not transduce signal output across the membrane by a piston-type displacement (40).

### Interpretation and assessment of the CG-MD results

To interpret the results of our GC-MD analysis, we categorised the signal output of these aromatically tuned EnvZ variants (40). For several two-component signalling circuits, including the EnvZ/OmpR, PhoQ/PhoP and CpxA/CpxR circuits, the steady-state output of the signalling circuits has been shown to be independent of the level of SHK present (40-43). However, in circuits containing the tuned variants of EnvZ, a different relationship between steady-state signal output and receptor level was observed, suggesting that the ratio of kinase to phosphatase activities was different within each receptor and always different than wild-type EnvZ (40). Based on this analysis, we found that the WLF-1 variant possessed the highest signal output while the WLF-5, WLF-4 and WLF-3 variants possessed slightly higher activity than wild-type EnvZ, which maintained receptor-concentration dependent robustness unlike the tuned variants. WLF-2 and WLF+1 were found to possess reduced signal output, while WLF+2 possessed the lowest overall signal output. These differences were previously quantified by analysing the slope of the change in CFP/YFP against the amount of receptor present and are visually represented in Figure S5 (40). These classifications are employed during interpretation of the CG-MD results in Figure 5.

**Figure 5.**
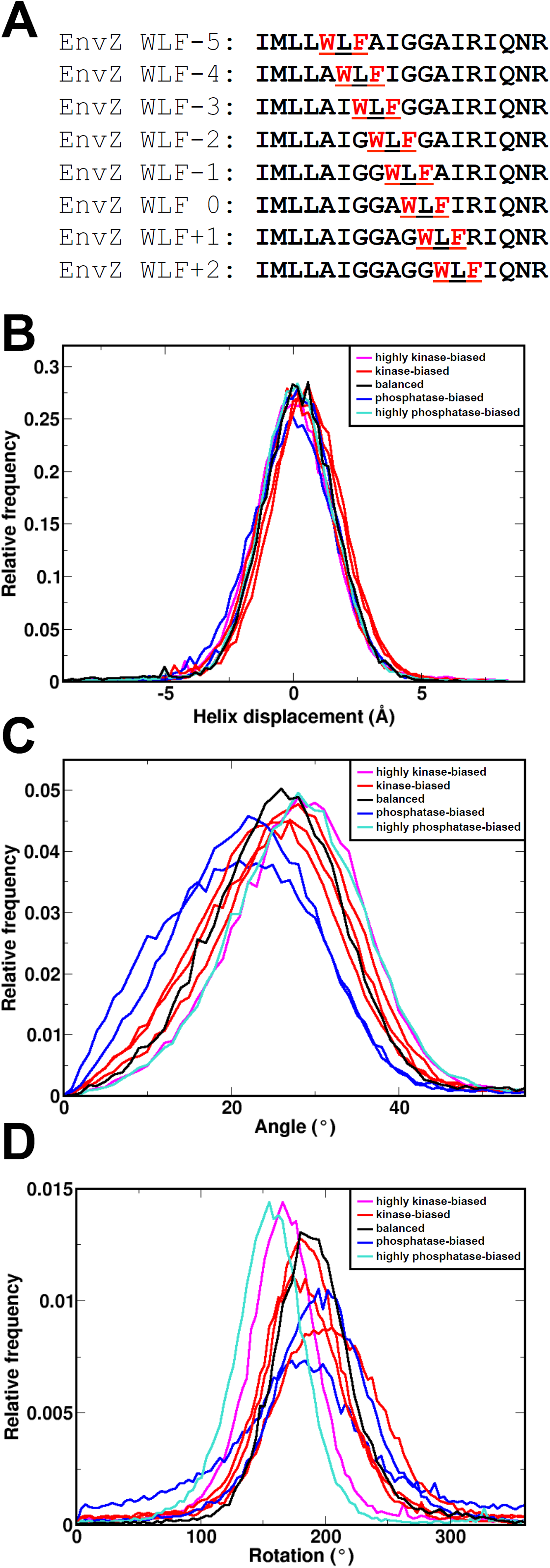
Aromatic tuning and signal output from EnvZ. (A) When aromatic tuning was performed in EnvZ, a Trp-Leu-Phe triplet (red) was repositioned within the C-terminal region of TM2. (B) Helix displacements, (C) tilt distributions and (D) azimuthal rotational distributions of the aromatically tuned EnvZ TM2 helices. Histograms are shown for all time points of all membranes of each ensemble. Aromatically tuned mutants have been classified based on their signal output as shown in Fig. S6 into five categories: highly kinase biased (WLF-1; pink), moderately kinase biased (WLF-5, WLF-4, WLF-3; red), balanced (wild-type; black), moderately phosphatase biased (WLF-2, WLF+1, dark blue) and highly phosphatase biased (WLF+2, bright blue).

Based on these previous results and the absence of asymmetric TM2 displacement observed within the *in vivo* sulfhydryl-reactivity assay (Fig. 4), we assessed whether molecular simulations would lend credibility to a non-piston type of transmembrane communication employed by EnvZ. We began by designing tuned TM2 sequences to subject to GC-MD simulations that matched those previously used during the aromatic tuning experimentation (Fig. 5A). Snapshots of individual frames and a movie from the CG-MD analysis with the wild-type EnvZ TM2 segment can be found in the Supporting Information (Fig. S6 and Movie S1). Unlike analogous experimentation with TM2 of Tar, no trend in helix displacement was observed (Fig. 5B, Table S1). This indicates that the mutations do not in isolation move the helix up and down in the membrane and would appear to rule out a pure piston motion as previously observed in Tar (35).

An alternative mechanism for transmembrane communication is to induce a tilt in the helix relative to the bilayer normal. This was originally proposed in Tar on the basis of crystallographic analysis of the receptor domain (44), and led to the proposal of a “swinging piston” where both displacement and tilt contributed to signal transduction. In addition, this would be consistent with a scissor-type model of TM communication proposed to be utilised by PhoQ (45, 46), which possesses a periplasmic PDC/CACHE domain (47). In testing the tuned EnvZ TM2 helices, a change in the extent of tilting was observed during the CG-MD simulations (Fig.5C, Table S2). It was found that a reduction in tilt, to a mean tilt of between 22 and 24 degrees, compared to the wild-type TM2 sequence with a tilt of approximately 26 degrees correlated with a decreased signal output (phosphatase-biased, dark blue). Interestingly, an increase in tilt to more than 29 degrees correlated with extremely biased signal output, *i. e.* kinase-dominant (WLF-1, bright red) and phosphatase-dominant (WLF+2, bright blue). Taken together, these results demonstrate that tilt, rather than vertical displacement, does correlate with modulated signal output from EnvZ.

Finally, a rotational/gearbox model of TMD-HAMP communication has also been proposed for EnvZ (48, 49). Central to this model is that interconversion between the kinase-dominant and phosphatase-dominant signalling states can be accomplished by rotation of individual HAMP helices by 26 degrees. With our simulations, we observed some consistency with this gearbox model of HAMP signalling, especially when the WLF-1 helix possessed a mean azimuthal rotation that is 25 degrees different than wild-type EnvZ (bright red in Figure 5D). In the full-length EnvZ WLF-1 receptor, this variant possessed the highest signal output. Further rotation of an additional 15 degrees for a total of 40 degrees from wild-type correlated with the most phosphatase-dominant receptor, WLF+2. For the other simulated helices, it was not possible to differentiate them into separate classes.

## Discussion

### Non-piston transmembrane communication by EnvZ

EnvZ has been shown to allosterically process cytoplasmic changes in osmolarity and upon interaction with MzrA within the periplasmic. Here, our *in vivo* analysis demonstrated that only the periplasmic end of EnvZ TM2 undergoes a conformational transition upon cytoplasmic stimulus perception and suggests that the asymmetric piston-type displacement employed by Tar is not used by EnvZ. To our knowledge, this is the first example of a periplasmic end of TM2 being affected by a cytoplasmic stimulus observed within EnvZ. Various experimentation has also been performed with the aromatically tuned variants of TM2 from both Tar and EnvZ. Previously, a linear correlation was observed between the position of the aromatic residue in Tar TM2, the position of the helices *in vitro* and *in silico* and the signal output from each Tar receptor (33, 34, 40, 50). Here, *in silico* analysis of EnvZ TM2 demonstrates that such a linear correlation is absent and that EnvZ functions by a nonpiston mechanism in which both tilting and azimuthal rotation play a substantial role in modulation of signal output (Figure 5).

### Correlations between domain composition and transmembrane communicationalic

Comparisons of recently published *apo* and *holo* high-resolution (~1.9 Å) crystal structures of the *E. coli* nitrate sensor NarQ that contain the periplasmic, TM and HAMP domains reveal extensive structural rearrangements involving a piston-like motion of TM1 relative to TM2 of approximately 2.5 Å. These displacements result in a lever-like rotation of individual HAMP domains upon binding of cognate ligand (36). Based on these results, the authors posit that receptors containing a membrane-adjacent HAMP domain function by a piston-type displacement of TM helices while those that lack such domains transduce signal by rotation of TM helices. We previously postulated a related yet different categorization of signaling mechanisms also based on the domain structure of bacterial receptors (51). We proposed that receptors containing a periplasmic four-helix bundle transduce signal across the membrane by piston-type displacements and that the attached membrane-adjacent HAMP domains might possess one of a multitude of signaling mechanisms including a gearbox-type rotation (48), a dynamic bundle (28) or potentially other mechanisms (51). Differentiating between these classification systems will provide a theoretical framework for understanding domain-based intraprotein allosteric communication by bacterial receptors. A recent authoritative structural-based review of transmembrane communication by bacterial receptors addresses these different suggestions (52).

This results presented here also examines whether SHKs that possess membrane-adjacent HAMP domains function solely by piston-type displacements or whether other signaling mechanisms might be employed. The results here with EnvZ should be compared with previous findings from the aspartate chemoreceptor (Tar) and the recent NarQ structures (36, 53-55) as these three are ideal candidates for comparison because they all possess a membrane-adjacent HAMP domain, however, while Tar and NarQ possess a periplasmic four-helix bundle, EnvZ possesses a periplasmic PDC/CACHE domain (37, 38). The authors of the recent NarQ structures posit that the presence or absence of the membrane-adjacent HAMP domain may be the difference between receptors employing piston-type mechanisms of transmembrane communication as compared to other mechanisms (36). However, differences employed during transmembrane communication by the Tar and EnvZ TMDs observed here and previously strongly suggest that Tar and EnvZ possess different mechanism of TM communication even though both possess a membrane-adjacent HAMP domain. Our previous work analyzed AS1 helices from *E. coli* NarX, *E. coli* Tar, *E. coli* EnvZ and Af1503, the HAMP domain resulting in the initial high-resolution structure (48), and found that the Tar and NarX AS1 helices possess similar properties, which the AS1 helices from both EnvZ and Af1503 fail to possess. Recent comparisons of the *apo* and ligand-bound structures of the combined periplasmic-TM-HAMP domain from *E. coli* NarQ demonstrate that binding of ligand results in symmetrical displacements of TM1 relative to TM2 of approximately 2.5 Å (36). These results are similar to Tar, which functions by asymmetrical TM2 displacement also possesses a periplasmic four-helix-bundle (39, 53-55).

### Conclusion

Based on these results, we conclude that *E. coli* EnvZ functions by a non-piston mechanism of transmembrane communication that is different than Tar, NarX and NarQ, which communicate across the membrane by piston-type displacements. Furthermore, we propose that TM signaling mechanisms can be predicted and assigned based upon the domain(s) present in the periplasmic region of a bacterial membrane-spanning receptor.

## Materials and Methods

### Bacterial strains and plasmids

*E. coli* strains DH10B (New England Biolabs) or MC1061 (56) were used for DNA manipulations, while strain K-12 MG1655 (57) served a non-fluorescent strain that was used to control for light scattering and cellular autofluorescence. *E. coli* strains MDG147 [MG1655 Φ(*ompF^+^-yfp^+^*) Φ(*ompC*^+^-*cfp^+^*)] (58) and EPB30 (MDG147 *envZ::kan*) (43) were employed for analysis of EnvZ signal output. As the C-terminus of bacterial receptors can be sensitive to the presence □of an epitope tag, we previously ensured that the addition of a V5-epitope tag did not alter the □signaling properties of EnvZ (40, 59). Plasmid pEB5 was employed as an empty control vector (41). Plasmid pRD400 (40) retains the IPTG-based induction of EnvZ from plasmid pEnvZ (60) while adding a seven-residue linker (GGSSAAG) (61) and a C-terminal V5 epitope tag (GKPIPNPLLGLDST) (62). Plasmid pEB5 was employed as an empty control vector.

### Selection of residues comprising TM2 of EnvZ

The primary sequence of EnvZ from *E. coli* K-12 MG1655 was subjected to DGpred using a minimal window of 9 residues and a maximal window of 40 residues (26). Alternatively, a software package that identifies TM helices with a Markov model (TMHMM v2.0) (27) was also employed. These software packages suggested that Leu-160 to Ile-181 and Leu-160 to Ile-179 comprise TM2 respectively. Based on these results and to maximize the probability of including all residues within TM2, we targeted all residues between positions 156 to 184 for the creation of a library of single-Cys-containing EnvZ receptors.

### Analysis of EnvZ signal output in vivo

Bacterial cultures were grown as described previously (40) with minor modification. MDG147 or EPB30 cells were transformed with pRD400 expressing one of the single-Cys-containing EnvZ receptors or pEB5 (empty). Fresh colonies were used to inoculate 2-ml overnight cultures of minimal medium A (63) supplemented with 0.2% glucose. Ampicillin, sucrose and IPTG were added as appropriate. Cells were grown overnight at 37 °C and diluted at least 1:1000 into 7 ml of fresh medium. Upon reaching an OD_600nm_ ≈ 0.3, chloramphenicol was added to a final concentration of 170 μg/ml. Fluorescent analysis was immediately conducted with 2 ml of culture and a Varian Cary Eclipse (Palo Alto, CA). CFP fluorescence was measured using an excitation wavelength of 434 nm and an emission wavelength of 477 nm, while YFP fluorescence was measured using an excitation wavelength of 505 nm and an emission wavelength of 527 nm. These values were corrected for cell density and for light scattering/cellular autofluorescence by subtracting the CFP and YFP fluorescence intensities determined for MG1655/pEB5 cells.

### Analysis of sulfhydryl-reactivity in vivo

Cells were grown as described above with minor chnges. Upon reaching an OD_600nm_ ~ 0.3, cells were subjected to between 250 μM molecular iodine for 10 min while incubating at 37 °C. The reaction was terminated with 8 mM N-ethylmaleimide (NEM) and 10 mM EDTA. Cells were harvested by centrifugation and resuspended in standard 6X non-reducing SDS-PAGE buffer supplemented with 12.5 mM NEM. Cell pellets were analysed on 10% SDS/acrylamide gels. Standard buffers and conditions were used for electrophoresis, immunoblotting and detection with enhanced chemiluminescence (64). Anti-V5 (Invitrogen) was the primary antibody and peroxidase-conjugated anti-mouse IgG (Sigma) was the secondary antibody. Digitized images were acquired with a ChemiDoc MP workstation (Bio-RAD), analysed with ImageJ v1.49 (65) and quantified with QtiPlot v0.9.8.10.

### CG-MD simulations with SIDEKICK

As previously described (35), coarse-grained molecular dynamics (CG-MD) simulations were performed using the MARTINI forcefield with an approximately 4:1 mapping of non-H atoms to CG particles. Lennard-Jones interactions between 4 classes of particles: polar (P), charged (Q), mixed polar/apolar (N) and hydrophobic apolar (C) were used to treat interparticle interactions. Within MARTINI, P and C particle types were subdivided to reflect varying degrees of polarity. Short range electrostatic interactions were treated Coulombically, shifted to zero between 0 and 12 Å. Lennard-Jones interactions were shifted to zero between 9 and 12 Å. α-helix integrity was maintained via dihedral restraints. Peptide termini were treated as uncharged. Simulations were performed using Gromacs 3 (66). Temperature was coupled using a Berendsen thermostat at 323 K (τ_T_□ = □1 ps), and pressure was coupled semi-isotropically (across XY/Z) at 1 bar (compressibility□ = □3×10^‒5^ bar^‒1^, τ_p_□=□10 ps). The initial simulation timestep was 20 fs. Initial models of the TM α-helices were generated as ideal, atomistically detailed a-helices using standard backbone angles and side-chain conformers. These were then converted to coarse-grained as described previously(35). Around 128 DPPC molecules were used in each simulation along with around 3000 CG water particles, giving a final system size of □65×65×13 Å.

## Acknowledgments

R. Y. was generously supported by the Indonesia Endowment Fund for Education, Ministry of Finance (S-4833/LPDP.3/2015). T. L. N. was supported by a grant from the Erasmus+ programme. R. R. D. was supported with start-up funding from the Faculty of Science and from the Institute of Biological and Biomedical Science (IBBS) at the University of Portsmouth. B. A. H. was supported by The Royal Society (UF130039).

## Supplemental Information

Figure S1. Steady-state expression of EnvZ variants containing a single Cys residue within TM2. EPB30/pRD400 cells expressing one of the single-Cys-containing variants were grown under the low-(0% sucrose) or high-osmolarity (15% sucrose) regimes. Under the low-osmolarity regime, reduced steady-state levels of the D156C variant were observed. In addition, disulfide formation was observed for the F157C and L160C variants in the absence of any additional oxidizing agent. When EPB30/pRD400 cells were grown under the high-osmolarity regime, the F157C, L160C and F161C variants exhibited disulfide formation in the absence of any oxidizing agent.

Figure S2. Steady-state signal output from the Cys-variant of EnvZ. (A) CFP and YFP fluorescence from MDG147/pEB5 (filled) and EPB30/pRD400 C277A (Cys-less; empty) cells grown under the low-(0% sucrose) and high-osmolarity (15% sucrose) regimes. (B) The CFP/YFP ratio from MDG147/pEB5 (filled) and EPB30/pRD400 C277A (Cys-less; empty) cells grown under the low-and high-osmolarity regimes estimates EmvZ signal output. Error bars represent standard error of the mean with a sample size of n ≥ 3. Reprinted from BBA Biomembranes, volume 1858, Annika Heininger, Rahmi Yusuf, Robert J. Lawrence and Roger R. Draheim, Identification of transmembrane helix 1 (TM1) surfaces important for EnvZ dimerisation and signal output, pages 1868-1875, Copyright 2016, with permission from Elsevier.

Figure S3. Signal output from the single-Cys-containing EnvZ variants. CFP (A) and YFP (B) fluorescence from EPB30/pRD400 cells expressing one of the receptors from the library grown under the low-osmolarity (0% sucrose) regime. CFP (C) and YFP (D) fluorescence from EPB30/pRD400 cells expressing one of the single-Cys-mutants from the library grown under the high-osmolarity (15% sucrose) regime. On the right axes, the ratio of signal output compared to EPB30/pRD400 cells expressing the Cys-less mutant is also presented to aid in comparison. In all panels, the shaded area represents the mean with a range of one standard deviation of the mean from EPB30/pRD400 cells expressing the Cys-less variant of EnvZ (Figure S2). Error bars represent standard error of the mean with an n 3.

Figure S4. Immunoblotting analysis of the sulfhydryl-reactivity experimentation. EPB30/pRD400 cells expressing one of the single-Cys-containing EnvZ receptors were grown under the low-(0%) or high-(15%) regimes until an OD_600nm_ of approximately 0.25 was reached. Cultures were then subjected to 250 μM molecular iodine. Particular Cys-containing EnvZ receptors resulted in the presence of dimeric EnvZ moieties that migrated at a slower rate than the monomeric species. A minimum of three immunoblots were used for each of the data points present in Figure 3.

Figure S5. Comparison of signal output from osmosensing circuits containing the various aromatically tuned EnvZ receptors. Steady-state signal output from circuits containing the WLF series of aromatically tuned variants expressed in EPB30/pRD400 cells grown under the low (upper panel) or high (lower panel) osmolarity regime is shown. The intracellular levels of phospho-OmpR are estimated through use of the antisymmetrical reporter system presented in Figure S1B. Signal output at low (open circles), medium (gray circles), and high levels (filled circles) of EnvZ-V5 expression (0.2, 0.5, and 0.8, respectively) is presented for comparison. The extent of sensitivity to changes in the amount of EnvZ present is also summarized as robustness. In this column, N/A represents not applicable, as in there is no reasonable amount of signal output, whereas REV indicates reversed, where a decrease in activity is observed as the level of EnvZ-V5 increases. Reprinted from ACS Synthetic Biology, volume 4, Morten N0holm, Gunnar von Heijne and Roger R. Draheim, Forcing the Issue: Aromatic Tuning Facilitates Stimulus-Independent Modulation of a Two-Component Signaling Circuit, pages 474-481, Copyright 2015, with permission from the American Chemical Society.

Figure S6. A randomly selected (number 63) simulation of the wild-type EnvZ TM2 was selected for demonstration. The blue and cyan dots represent water molecules, while the bronze spheres represent the phosphate in DPPC. The backbone carbon atoms of the peptide are shown pink, while the yellow spheres represent side chains.

Table S1: Mean displacement of the EnvZ TM2 segments in the membrane bilayer.

Table S2: Mean tilt of the EnvZ TM2 segments in the membrane bilayer.

Table S3: Mean azimuthal rotation of the EnvZ TM2 segments in the membrane bilayer.

Movie S1. A movie composed of frames from the simulation used for Figure S6. To compose the movie, 500 frames spaced 200 ns apart were selected. The blue and cyan dots represent water molecules, while the bronze spheres represent the phosphate in DPPC. The backbone carbon atoms of the peptide are shown pink, while the yellow spheres represent side chains.

